# Consensus clustering of herpesvirus protein interaction networks provides insights into their evolutionary relationship with bacteriophages

**DOI:** 10.1101/473751

**Authors:** Anna Hernández Durán, Todd M. Greco, Benjamin Vollmer, Ileana M. Cristea, Kay Grünewald, Maya Topf

**Affiliations:** Institute of Structural and Molecular Biology, Birkbeck College, University of London, Malet street, London, WC1E 7HX, UK; Division of Structural Biology, Wellcome Centre for Human Genetics, University of Oxford, Roosevelt Drive, Oxford, OX3 7BN, UK; Department of Molecular Biology, Princeton University, Lewis Thomas Laboratory, Washington Road, Princeton, NJ 08544, USA; Department of Structural Cell Biology of Viruses, Centre for Structural Systems Biology, Heinrich Pette Institute, Leibnitz Institute of Experimental Virology, University of Hamburg, Notkestr. 85, Hamburg, D-22607, Germany

## Abstract

Infections with human herpesviruses are ubiquitous and a public health concern worldwide. Current treatments reduce the morbidity of some manifested symptoms but neither remove the viral reservoir from the infected host nor protect from the recurrent symptom outbreaks that characterize herpetic infections. The difficulty in therapeutically tackling these viral systems stems in part from their remarkably large proteomes and the complex networks of physical and functional associations that they tailor. This study presents our efforts to unravel the complexity of the interactome of herpes simplex virus type 1 (HSV1), the prototypical herpesvirus species. Inspired by our previous work, we present an improved computational pipeline for the protein-protein interaction (PPI) network reconstruction in HSV1, combining both experimentally supported and bioinformatic predictions. Our newly-developed consensus clustering approach allows us to extend the analysis beyond binary physical interactions and reveals higher order functional associations including that of pUS10 with capsid proteins. In-depth bioinformatics sequence analysis unraveled structural features of this 34-36 kDa protein reminiscent of those observed in some capsid-associated proteins in tailed bacteriophages, with which herpesviruses are thought to share a common ancestry. This suggests that pUS10 could represent an evolutionary vestige between these two viral lineages. Using immunoaffinity purification-mass spectrometry we found that pUS10 specifically co-isolated with the inner tegument protein pUL37, which binds cytosolic capsids, contributing to initial tegumentation and eventual virion maturation. In summary, this study unveils new insights at both the system and molecular levels that can help better understand the complexity behind herpesvirus infections.

## Introduction

Herpesviruses infect a wide range of eukaryotic organisms, and are the etiologic agent of severe diseases in livestock and humans. At present, there are nine species of herpesviruses known to routinely infect human. They are referred to as the human herpesviruses, and their infections are associated with symptoms ranging from fever and cutaneous lesions, to encephalitis, meningitis, and a number of cancerous malignancies [1]. A link between herpetic infections and the neurodegenerative Alzheimer’s disease was recently confirmed, emphasising the socio-economic burden associated with these viruses [2,3].

Herpesviruses are enveloped viruses that assemble into a morphologically unique extracellular particle (i.e. virion), which is organised in concentrical structural layers [4]. At the innermost of the virion particle, an icosahedral protein shell of ~ 120nm diameter - the capsid - encloses the double-stranded DNA (dsDNA) genome of the virus. Surrounding the capsid, a protein matrix - the tegument - occupies about two thirds of the virion particle volume. The tegument contains proteins that are delivered into the host cytoplasm upon infection and therefore are ready to initiate their function prior to the transcription of any viral genes. The entire particle is coated by a lipid bilayer - the envelope - that contains several proteins and glycoproteins that are crucial for host cell entry and cell-to-cell spread, as well as for modulation of the host’s immune response. As for other enveloped viruses, entry into the host cell occurs via fusion of the envelope with plasma membrane or endocytic cellular membranes, which results in release of the virion content into the host cell cytosol [5]. Altogether, herpesviruses are composed of between ~70 to ~170 different protein species and achieve diameters between ~150 to ~250nm [6,7].

To understand the behavior of a system, the relationships among the components are as important as the components themselves. The need for a comprehensive description of the relationships among viral proteins, and with host factors, has prompted numerous protein-protein interaction (PPI) studies. In 2016 we published a compilation of PPI data in herpes simplex virus type 1 (HSV1), the prototypical species of the *Herpesviridae* family based on integration of experimentally supported data from five public resources [8]. Here, we present an improved computational pipeline for PPI network reconstruction, collating data from seven public repositories with computationally predicted PPIs inferred using conservative sequence similarity assessments. The reliability of all gathered PPIs is measured using a new scoring scheme. Moreover, we applied network analysis approaches to gain insights into the functional organization of the extracellular viral particle. We developed a clustering consensus protocol to study the community structure underlying the binary interactions among virion proteins. The resulting higher order relationships predicted previously unrecognized functional viral protein associations, such as between the inner tegument protein pUL37 and pUS10. We further investigated this relationship using bioinformatic sequence analysis, and provided experimental support using immunoaffinity purification-mass spectrometry in primary human fibroblasts infected with HSV1.

## Results

### PPI network assembly

Binary Protein-Protein interaction (PPI) data were obtained from five molecular interaction repositories (BioGRID [9], the Database of Interacting Proteins (DIP) [10], IntAct [11], Mentha [12], and VirHostNet 2.0 [13]) and two structural databases (Protein Data Bank (PDB) [14] and the Electron Microscopy Data Bank (EMDB) [15]) (Fig 1A). We collected PPIs that have been detected for all nine human herpesvirus species, and three closely-related non-human herpesviruses (Fig 1B and S1 and S2 Tables). The latter were included as these species are frequently used as animal models of herpetic human infections (see Materials and Methods). The resulting joined non-redundant data set contained 2854 unique pieces of evidence, 2363 unique PPIs, and 758 unique protein sequences. From this data set, PPIs experimentally detected in HSV1 were directly added to the network (Fig 1C). PPIs experimentally detected in species other than HSV1 were used to predict new PPIs in HSV1 (Fig 1D). To predict PPIs, we used an orthology-based method referred to as *interologues mapping* [16]. This method predicts an interaction in HSV1, if both putative interactors have homologues known to interact in other species (see Materials and Methods). Homology relationships are inferred based on sequence-based Hidden Markov Model (HMM) profile alignments [17] in combination with a conservative multi-criteria threshold to filter out potential spurious matches from the homology search results (see Materials and Methods).

**Fig 1.**
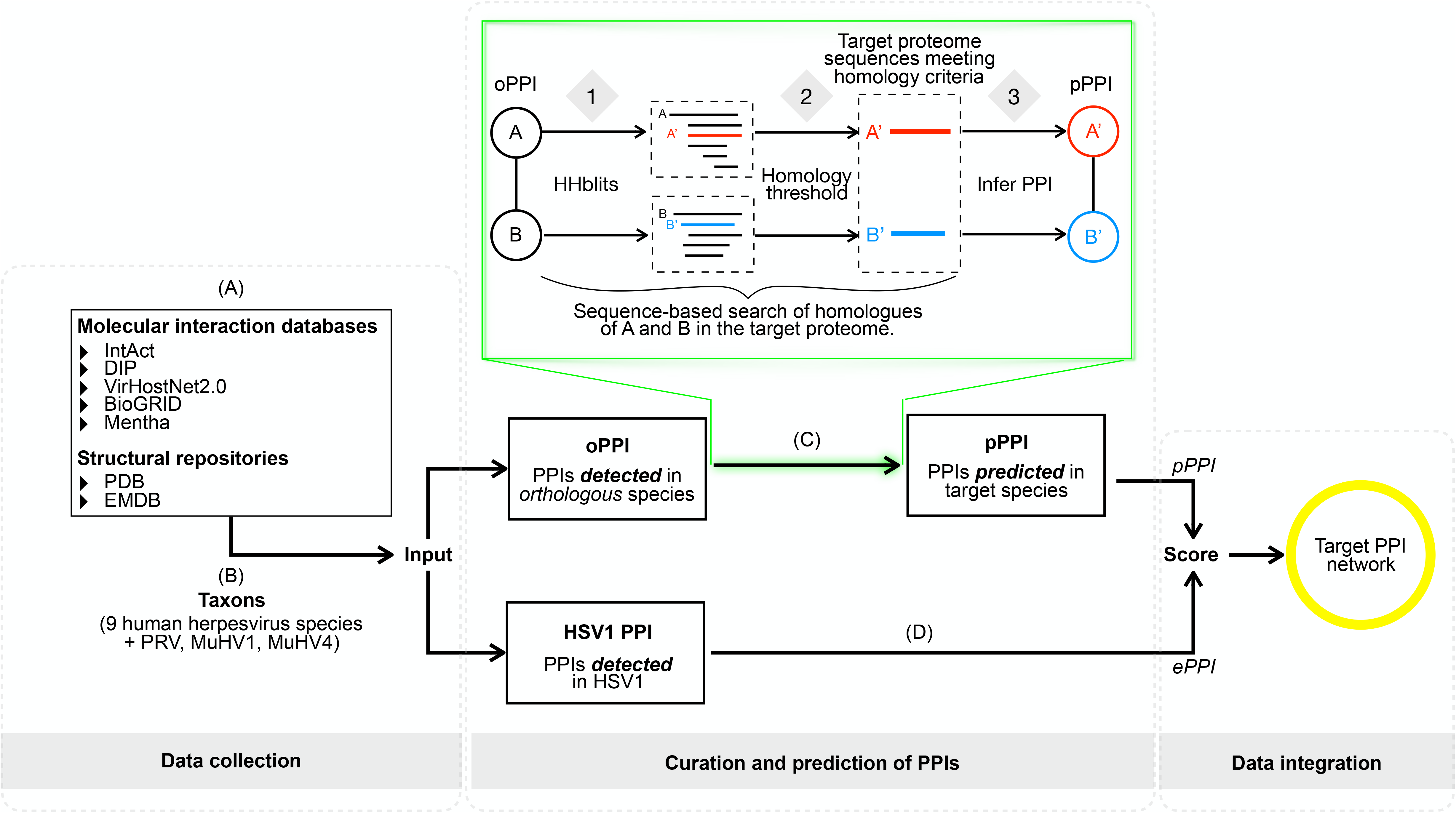
Network assembly framework. (A and B) PPI data for a total of twelve herpesvirus species (nine human and three non-human herpesviruses, together covering members of all three subfamilies, i.e. the α-, β- and γ- *Herpesvirinae*, S1 and S2 Tables) were collected from seven public resources [9–15]. (C) PPIs experimentally detected in HSV1 were transferred to its interactome (ePPIs). (D) PPIs detected in any of its orthologous herpesvirus species (oPPIs) were used to predict PPIs in HSV1 (pPPIs). Predictions were conducted based on a sequence-based interologues mapping [16] approach (green box), and included the following steps: for each protein involved in a binary oPPI, (1) sequence-based homologous sequences in the HSV1 proteome were searched for using HHblits [17]; (2) a conservative homology threshold was applied to filter out potential spurious matches among the list of candidates returned by HHblits. From the remaining matches, the best scoring sequence was selected as the most reliable putative HSV1 homologue; (3) if potential HSV1 homologous sequences were found for both proteins in the initial oPPI, an interaction between the two HSV1 sequences was predicted. (E) Predicted and experimentally supported PPIs were joined into a non-redundant data set, and scored based on their supporting evidence (see Materials and Methods).

Experimentally detected interactions in HSV1 and computational predictions were compiled into a PPI network, and the confidence of each interaction based on its cumulative evidence was assessed using a new scoring function (see Materials and Methods). The new function is a modification of the MIscore function [18], which was developed by the Proteomics Standards Initiative [19], and is compliant with standardised protocols for the assessment and representation of molecular interaction data [20]. Our function essentially adds a conservative penalty function term to the original MIscore function for interactions without experimental support. This term includes information on the prediction method (in this case sequence-based alignment) and the total number of homologous species from which the prediction was inferred. The function gives higher confidence scores to interactions that have been predicted from a larger number of species. For instance, if an interaction can be predicted in HSV1 based on interactions experimentally detected in three other species of herpesviruses (e.g. VZV, EBV and HCMV), the prediction will score higher than if it had been obtained from a single experimental observation (e.g. only in VZV).

The reconstructed network (Fig 2 and S3 Table) contained 369 PPIs, formed among 68 proteins, and supported by 643 unique pieces of evidence. Among the 369 PPIs, there were 159 experimentally supported interactions, and 250 computationally predicted; 40 interactions were supported by both experimental and computational evidence. All these network data are available through our new version of the HVint database [8]: http://topf-group.ismb.lon.ac.uk/hvint/.

**Fig 2.**
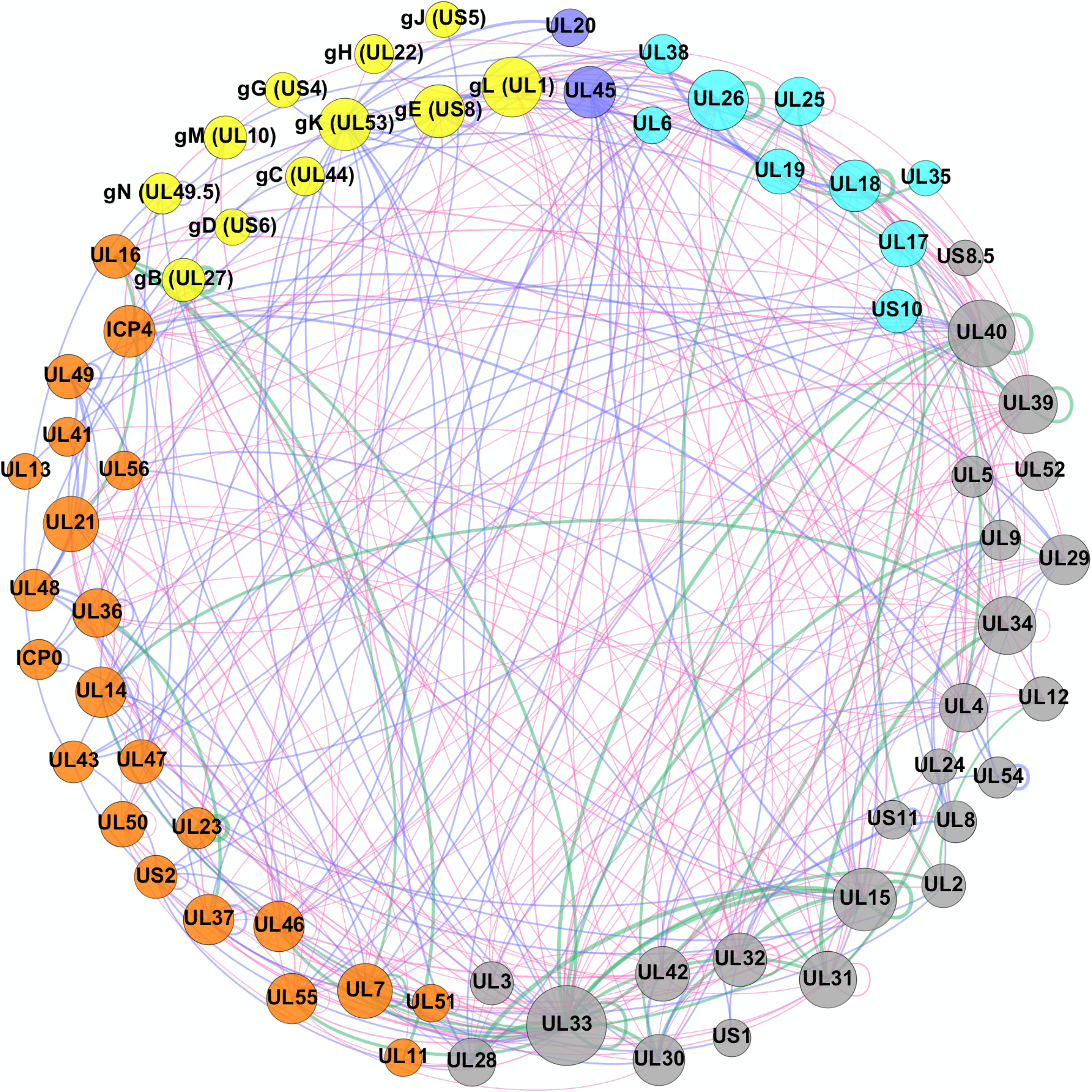
Reconstructed PPI network for HSV1. Nodes represent proteins and their size is proportional to the number of interactions associated with each protein in the network (degree). Nodes are colour-coded as follows: cyan for capsid and capsid-associated proteins, orange for tegument proteins, yellow for envelope glycoproteins, blue for non-glycosylated envelope proteins, respectively, and grey for proteins that are not present in the mature virion particle (i.e. typically only expressed during intra-cellular stages). Edge (or link) thickness reflects the confidence score for the interaction (the thicker, the higher the confidence). Edges are colour-coded to indicate the type of supporting evidence behind it, i.e. blue for experimentally supported interactions, red for computationally predicted, and green for interactions with both experimental and computational supporting evidence.

### Selection of high confidence PPI predictions for experimental testing

We assessed the ability of our protocol for predicting meaningful interactions for experimental testing. We conducted a manual literature review in search of experimental support for our best predictions (5% top scoring interactions) among published literature that had not been included in the input dataset (i.e. it was not used to infer the predictions). This search returned experimental support for 4 out of the total 11 (top 5% scoring) predicted PPIs examined [8,21,22] (Table 1).

**Table 1.**
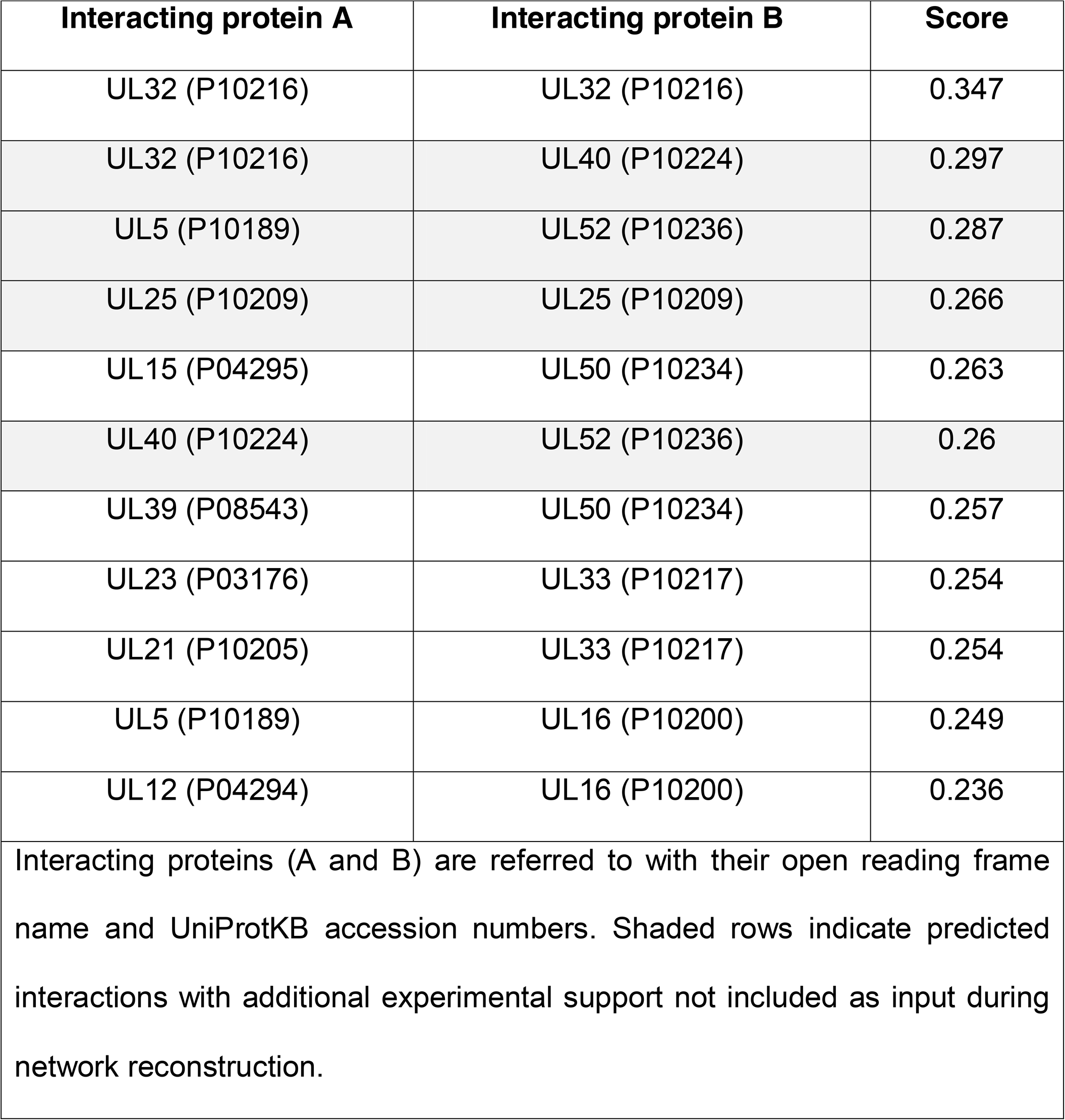
Top 5% computationally predicted PPIs.

### Analysis of the community structure in the HSV1 network

Using the HSV1 network data compiled at this point, we undertook the analysis of the community structure of the subnetwork formed by virion proteins. These are defined as components of the extracellular viral particle (i.e. capsid, tegument, and envelope proteins), which are currently well identified in this species [23–26]. We applied two further steps to remove noise (e.g. false positive PPIs) in the network to reduce potential sources of artefacts in the clustering results (Fig 3A, see Materials and Methods). First, we removed PPIs in the network taking place between capsid (and capsid-associated) and envelope proteins, which are likely unfeasible due to spatial constraints imposed by the virion tegument. Second, we extracted, from the network obtained in the previous step, the largest <u>c</u>onnected <u>c</u>omponent (LCC – the largest group of nodes in which any pair of them can be connected by a path). The resulting network contained 40 nodes and 94 edges.

**Fig 3.**
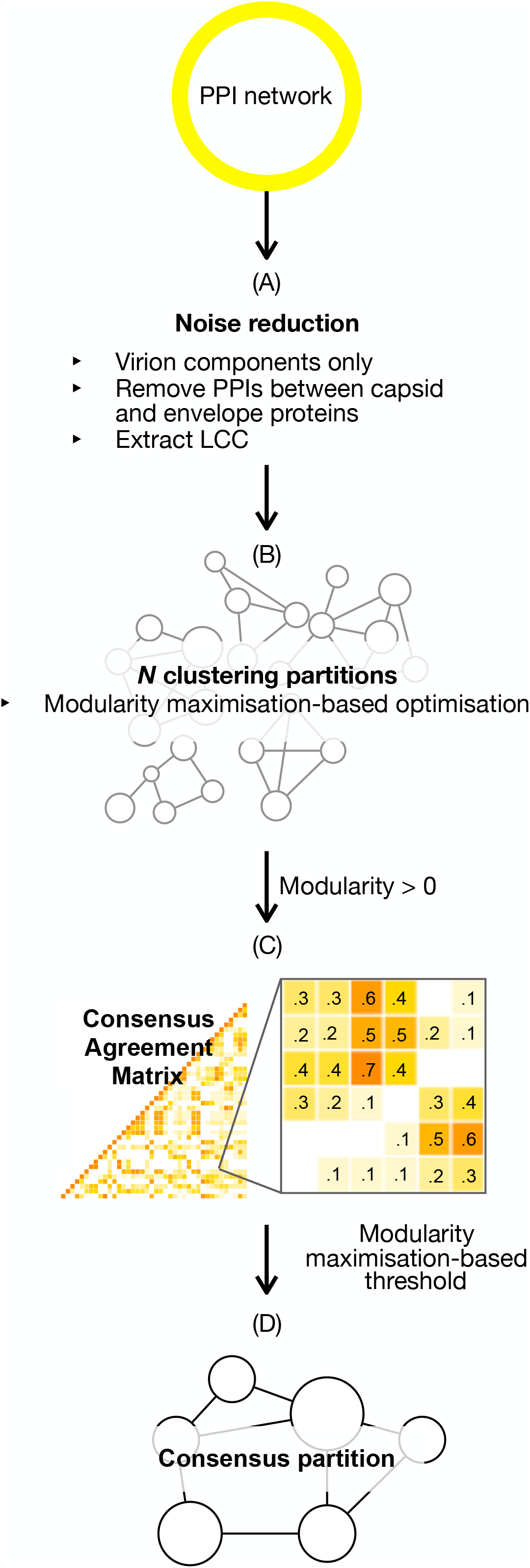
Consensus clustering protocol. Starting from an input network, (A) a series of steps were applied to remove potential false positives. Here, these steps included selection of virion components only, and removal of PPI between capsid (and capsid-associated) proteins to envelope proteins. (B) N=14 clustering algorithms were applied to the network. (C) The output base partitions with positive modularity were integrated into a consensus agreement matrix. (D) A final consensus clustering partition was derived by filtering out the values in the consensus agreement matrix based on a modularity-maximisation threshold.

Next, 14 different clustering algorithms were applied to this LCC network (Fig 3B and S4 Table). Those output partitions yielding positive modularity (11 in total) were integrated into a consensus agreement matrix (Fig 3C). Values in the consensus agreement matrix cells represented, for each pair of proteins in the network, the fraction of partitions that clustered the pair together. Using a threshold-based method guided by modularity maximization a consensus partition was inferred from the consensus agreement matrix (Fig 3D). This partition achieved a modularity of 0.43 (for thresholds in [0.55, 0.60]) (S1 Fig and S5 Table); values between 0.3 and 0.7 are considered indicative of a significant community structure [27,28].

The consensus partition divided the network into four communities (Fig 4). Their biological consistency was then assessed using functional annotation data manually curated from Gene Ontology (GO) [29,30] annotations and published literature.

**Fig 4.**
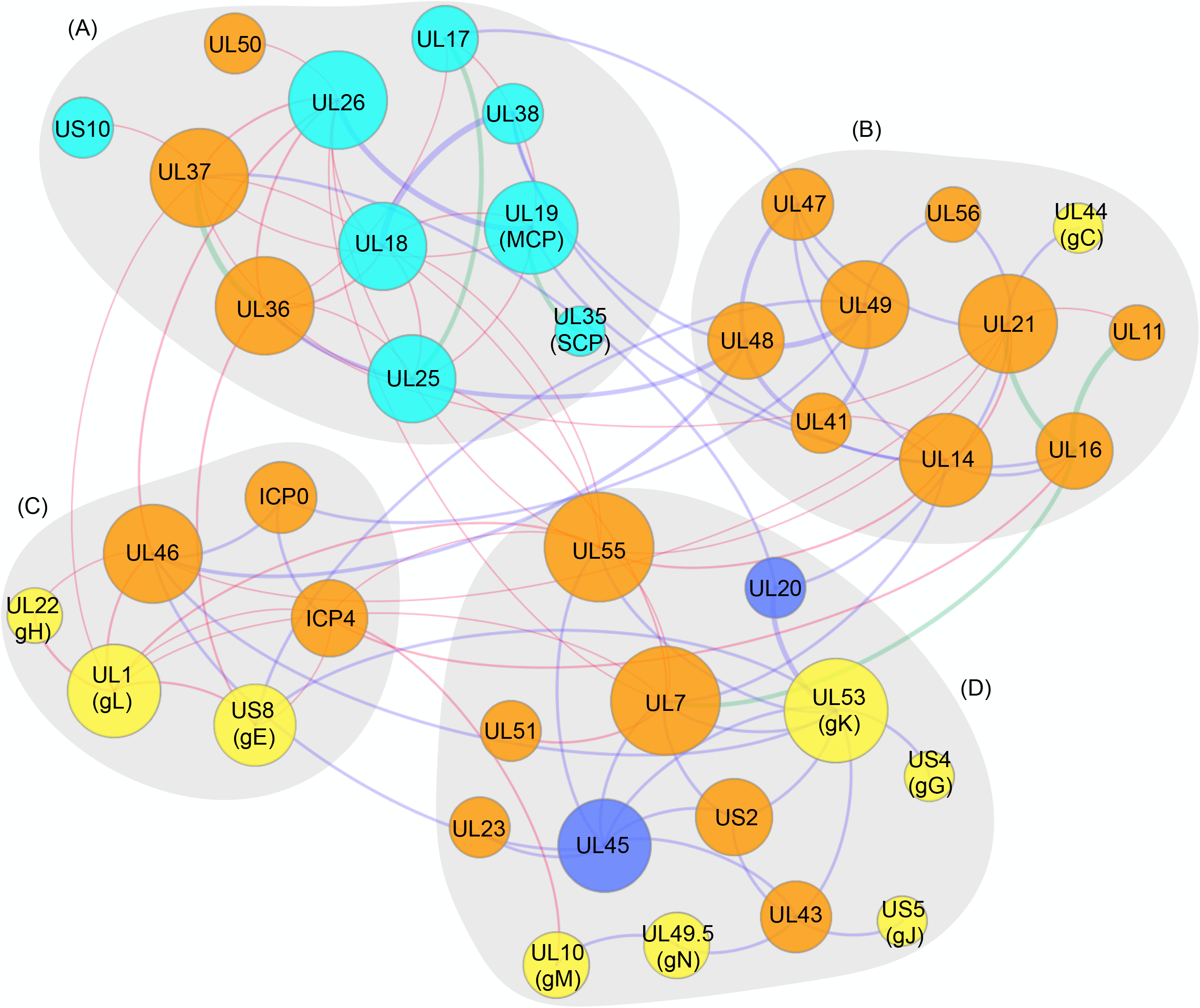
Community structure inferred for the HSV1 virion network. Depiction of nodes and edges follows the same criteria as described in Fig 2. Four separate communities are indicated by grey shaded areas.

#### Community A

Community A illustrates almost entirely the composition of the capsid (Fig 4). The only capsid protein missing is pUL6 (the portal protein), which appears disconnected from the LCC in the virion subnetwork. This was surprising as the localisation of pUL6 in the capsid, forming the capsid portal complex, is well established [31]. However, the input data sets used to feed the network assembly framework did not contain evidence for this interaction.

Community A also contains the protein pUS10. pUS10 is a poorly characterised α-subfamily specific protein of 312 residues, which has been found in the nuclear, peri-nuclear and cytoplasmic cellular compartments and co-precipitating with capsids, yet direct interactions with capsid proteins have not been reported. It exists in two phosphorylation states and is regarded as a minor component of the tegument [32]. A consensus zinc finger was identified in pUS10 homologues although the protein failed to bind nucleic acids (common among zinc finger proteins) during experimental testing [32,33]. Finally, HSV1 pUS10 presents a high proline content at its N-terminus and a four-residue long polyproline sequence located centrally. To assess whether the clustering of pUS10 with capsid proteins had a functional significance or it was an artefact of a low number of interactions for pUS10 in the input graph, we sought further characterization of the protein through primary sequence analysis (Fig 5 and S2 Fig, see Materials and Methods). An initial search for potential sequence homologues did not identify any candidates beyond pUS10 counterparts, yet it highlighted the presence of seven tandem collagen-like repeats (CLRs) located toward the N-terminus of the protein sequence. Interestingly, through a manual curation of the literature, we found support for this prediction in a 30-year old study [34] (note that pUS10 was annotated by its MW in this study, which could explain why this prediction was not included in databases such as UniProtKB). This prominent feature prompted compelling hypothesis on its functions and evolutionary history (see Discussion). Secondary structure predictions indicated that the N- and C-termini of the protein are structurally distinct. Whilst the N-terminus was predicted to be disordered, the C-terminus was rich in α-helices. Additionally, our predictions revealed a potential single-pass transmembrane segment at the very C-terminus of the protein. This prediction overlaps with the potential zinc finger motif, which could explain why so far no functional evidence has been provided.

**Fig 5.**
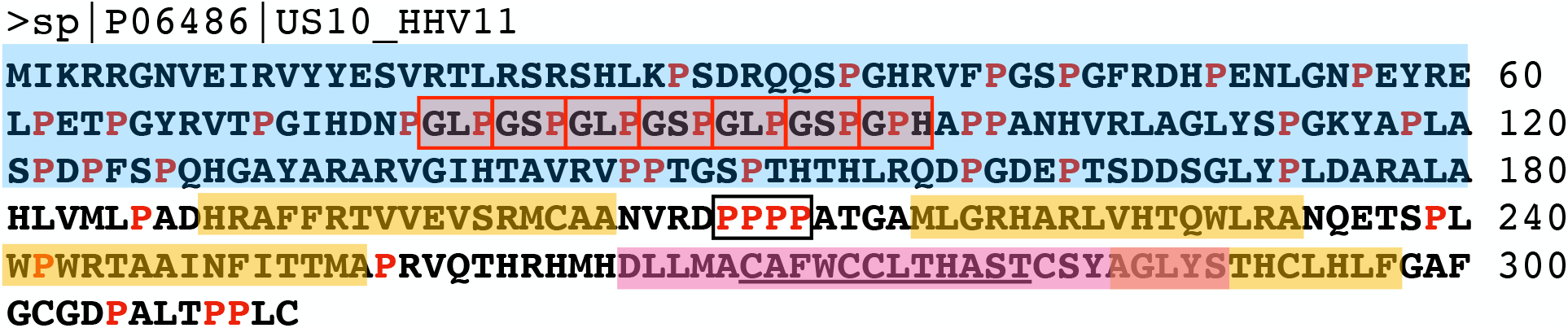
Sequence characterization of pUS10. Both previously described and newly identified features are indicated. Predicted disordered regions, α-helices and transmembrane helices are highlighted in blue, yellow and pink, respectively. Collagen-like repeats (CLRs) are framed in red boxes. Prolines are highlighted in red. The four-residues polyproline sequence is framed in a black box. The previously found consensus zinc finger sequence [33] is underscored.

The binary interaction that connects pUS10 with other components within its community is the interaction with pUL37. This interaction was predicted by our orthology-based method, and we aimed to further validate it experimentally (see below). pUL37 HSV1 is an essential inner tegument protein [35] that contains a binding site for HSV1 pUL36, through which it is recruited to cytosolic capsids before secondary envelopment [36,37] or to sites of secondary envelopment in the absence of capsids [38].

### Experimental support of the pUS10-UL37 interaction by immunoaffinity purification – mass spectrometry

To provide evidence in support of the predicted pUS10-UL37 interaction, we isolated pUL37 from infected cells using immunoaffinity purification and analysed the co-isolated proteins by quantitative mass spectrometry (IP-MS). Specifically, protein complexes were isolated from human fibroblasts synchronously infected with HSV1 strains encoding either pUL37 tagged with enhanced green fluorescent protein (pUL37-EGFP) or EGFP alone as a control. Isolations were performed at two functionally distinct time points of infection, 8 and 20 hours post infection (hpi) (S6 Table). Eight hpi represents an early time point of pUL37 expression, which is prior to secondary envelopment. This is consistent with our observation that pUL37-EGFP appears diffusely localized in the cytoplasm by epifluorescence microscopy (Fig 6A). In contrast, at 20 hpi secondary envelopment and virus particle release is in progress, reflected by pUL37-EGFP fluorescence visualized as capsid-associated puncta within the cytoplasm and maturing virions (Fig 6A). The observed temporal kinetics and localization of pUL37 induction were consistent with the prior characterization of this pUL37-EGFP HSV1 strain [37].

**Fig 6.**
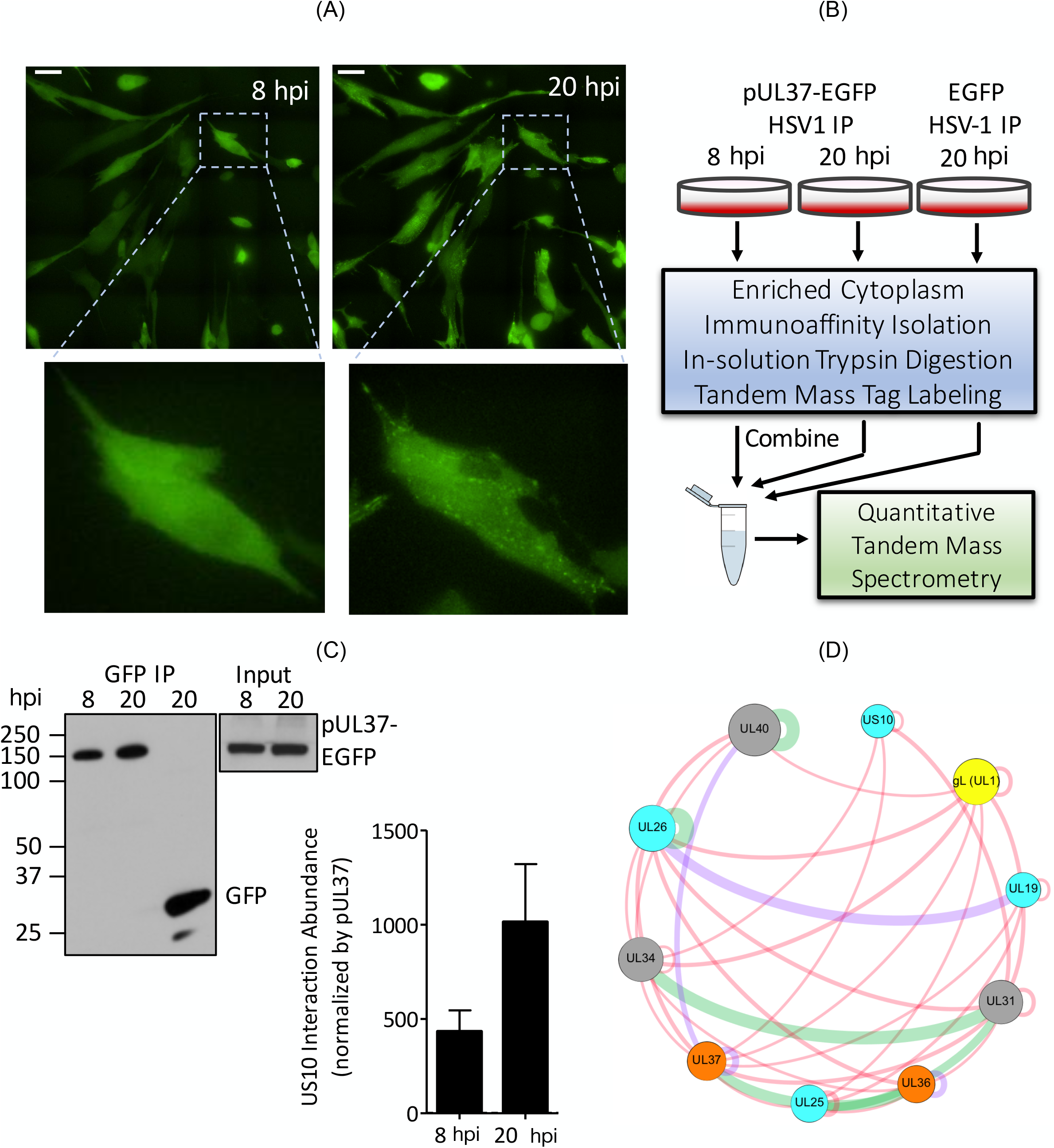
Experimental validation of the pUL37-US10 interaction in HSV-1 infected primary human fibroblasts. (A) Visualization of pUL37-EGFP during HSV1 infection of human foreskin fibroblasts using live cell epifluorescence microscopy. Images show a representative field of infected cells at 8 and 20 hpi. Zoomed images show localization of pUL37-EGFP (green) in the same cell at 8 and 20 hpi. Scale bar = 50 µm. (B) IP-MS workflow. Human fibroblasts were synchronously infected (multiplicity of infection = 10) with either pUL37-EGFP or EGFP HSV1, with two replicates per condition. HSV1-UL37GFP was collected at 8 and 20 hpi and HSV1-GFP at 20 hpi (HSV1-GFP). pUL37-EGFP and its interactions were isolated from the cytoplasmic cell fraction by immunoaffinity purification using anti-GFP antibodies. Proteins were digested with trypsin, and the resulting peptides from each sample were labelled with unique TMT reagents and then combined prior to nanoliquid chromatography-tandem mass spectrometry analysis. (C) The recovery of pUL37-EGFP and EGFP in the immunoisolates was assessed by western blot with an anti-GFP antibody. 10% of each sample was analysed. (D) PPIs around pUL37 present in the reconstructed HSV1 network (Fig 2) and supported by immunoaffinity purification results. (E) The abundance of pUS10 interaction with pUL37 at 8 and 20 hpi (average ± range, N=2). The relative amount of pUS10 was calculated by TMT-MS quantification and normalized by the respective pUL37 TMT abundance in each IP.

After establishing the temporal expression and cellular distribution of pUL37, we performed IP-MS analyses of pUL37-EGFP and EGFP complexes isolated from cytoplasmic-enriched lysates (Fig 6B). The isolation of pUL37-EGFP from the input lysates was confirmed by western blot (Fig 6C). Next, the co-isolated proteins were identified and relatively quantified using labelling with isobaric with Tandem Mass Tags (TMT). Specifically, proteins were digested with trypsin, and the resulting peptides were derivatised with distinct TMT labels. This approach offered multiplexing, as the labelled pUL37-EGFP and EGFP IPs performed in biological replicates (N = 2 replicates) were combined and analysed by tandem MS (Fig 6B). Therefore, both the specificity of the pUL37 interactions (via comparison to the control EGFP IP) and the relative abundance of the interactions at the different time points of infection (8 and 20 hpi) could be simultaneously assessed in this quantitative MS workflow. Nine viral proteins, including pUS10, that appear as interacting with pUL37 in our network were quantitatively enriched by ≥2-fold in pUL37-EGFP vs EGFP control IPs in at least one-time point and in both replicate experiments (S6 Table). After normalizing the pUS10 abundance to the abundance of pUL37-EGFP bait calculated by TMT-MS, we observed that the relative amount of pUS10 co-isolated with pUL37 was increased at the later stage of infection. At this stage, pUL37-EGFP is localized to sites of secondary envelopment and associates with maturing virions, consistent with the formation of pUL37-EGFP puncta at 20 hpi (Fig 6A). These data suggest a potential role for the pUL37-US10 interaction in the cytoplasm during virion maturation.

#### Community B

In community B (Fig 4), proteins pUL41, pUL47, pUL48, and pUL49 are associated, directly or indirectly, and are all known to modulate gene expression [39–41]. Another major group of proteins, (pUL11, pUL14, pUL16, pUL21, and pUL6) are related to early virion morphogenesis, i.e. recruitment of capsids and virion components to the enveloping membranes during early stages of tegumentation [42–45]. The tight relationships among the gene expression modulation proteins have been well characterized, yet the steps that guide their incorporation into the virion particle are not yet clearly defined. The results of our clustering analysis suggest that this process could be guided by the coordinated action of cluster B members involved in virion morphogenesis.

#### Community C

Community C is defined by proteins involved in late stages of secondary envelopment and virion release. This community is enriched in envelope glycoproteins (gM/gN, gK/UL20, pUL45, pUL43, gJ, and gG) that regulate membrane-associated events. These include internalization of proteins from the plasma membrane, membrane fusion rates, and modulation of immune responses. Tegument proteins in this community are also annotated with the trafficking of virion components at late stages of virion morphogenesis [46–52].

An interesting observation from community C comes from the presence of pUL55, another poorly characterised α-subfamily-specific protein [53]. Our primary sequence analysis of this 186-residue protein failed to find characteristic structural features but predicted pUL55 to be an α/β protein. The analysis of the hierarchical structure of community C on the other hand (S3 Fig, see Materials and Methods), did suggest a very stable clustering tendency with four other proteins in the cluster, namely gK/UL20, pUL45, and pUS2. The cluster assignment together with the particularly strong association with the latter four proteins [54–57] suggest new functional scenarios in late envelopment and virion release events for the currently poorly-understood pUL55 protein.

#### Community D

Community D is the most functionally inconclusive community. We hypothesise that this is due to the lack of known binding partners (either due to missing data in the input data sets or to failed homology mappings) in the network. Specifically, the lack of proteins gI and pUS9, both associated to gE (gI physically and pUS9 functionally [58,59], as well as proteins gB and gD, which are the members of the fusion machinery (together with gH/gL [5]).

## Discussion

Human herpesviruses are ubiquitous human pathogens with a severe socioeconomic impact worldwide [1]. Key to a successful infection is the delicate coordination of the complex network of virus-virus and virus-host protein interactions. Human herpesviruses encode distinctively large proteomes, which translate into large numbers of possible intra-viral PPIs. Several studies have attempted to address the description of these networks using a variety of, mostly, experimental approaches [60–63].

The aim of this study was to undertake a systematic and comprehensive analysis of PPI data in HSV1, the prototypical species of human herpesviruses, to characterize the binary and higher order relationships among its encoded proteins. We addressed this using computational approaches to integrate and complement experimental binary PPI data, and analyse its modular structure.

A first step was to design a new computational framework for reconstructing the PPI network. The three pillars of this framework are: a) data integration; b) standardization; and c) reproducibility. Our method integrates experimentally supported and computationally predicted binary PPI data in a non-redundant manner using standardized molecular interaction data formats. Sequence-based orthology relationships are the basis of our PPI prediction approach. The search for homologues is conducted using the HHblits algorithm, the sensitivity and selectivity of which currently outperforms other available methods [17]. Next, our method applies more conservative criteria than previously [8] to select only the most confident candidate homologues among the results returned by HHblits. Additionally, all interactions are assessed under a common scoring scheme inspired by the standardized MIscore function [18], and which incorporates a new term to penalise the scores of computational predictions in a non-linear way. This term applies to interactions predicted based on protein conservation, and includes information on the prediction method (here, sequence-based interologues mapping), and the number of species from which the interaction is predicted. This aims to reflect that a broader cross-species conservation of protein interactions is indicative of a greater functional relevance of the conservation of the interaction. As a result of our updated, more conservative, protocol, we observe that although the initial PPI dataset is larger than that used in our previous study (due to additional input databases) [8], the final PPI network is smaller. The reconstructed network is freely available to explore through the newly updated HVint database [8] interface: http://topf-group.ismb.lon.ac.uk/hvint/.

We tested our predictive strategy for the top 5% predicted interactions and we were able to find independent experimental support (i.e. not used to build the network) for 4 out of these 11 interactions. These included PPIs between pUL40 and pUL32, pUL40 and pUL52, pUL5 and pUL52, and the homodimer formed by pUL25 [8,21,22]. This highlights the power of our PPI prediction protocol in proposing new interactions for testing.

The community structure analysis performed on the virion subnetwork of HSV1 indicated the presence of four large communities, consistent with distinct stages of the virion formation process. This analysis suggests new functional relationships among virion components. The delineated communities illustrate events related to capsid formation, early tegumentation at peri- and juxtanuclear regions, and late tegumentation and virion release steps taking place in TGN-derived vesicles and the plasma membrane, respectively.

Community A drew our attention to protein pUS10, so far, a poorly characterized minor component of the virion specific to the α-*Herpesvirinae* subfamily. Our primary sequence analysis of pUS10 predicts the sequence to be structurally divided into a disordered N-terminus, and an α-helical C-terminus likely to embed a single-pass transmembrane segment. Additionally, our analysis identified a seven-repeat length CLR, centrally in the protein sequence. CLR sequences are characterized by the GXY pattern, where X and Y are any amino acid, and adopt left-handed helical conformations that tend to tightly pack into trimeric right-handed helices [64,65]. CLRs have been identified in a range of organisms, from multicellular eukaryotes to unicellular bacteria and viruses, and they participate in a wide range of processes including e.g. adhesion, morphogenesis, and regulation of signaling cascades among others [64–67]. These features became especially interesting as they are highly reminiscent to those observed in another CLR-containing protein, gp12, from the bacteriophage SPP1 [68,69]. Importantly, SPP1 is one of the species of tailed dsDNA bacteriophages that have been shown to share a common evolutionary ancestry with herpesviruses [70]. Protein gp12 from SPP1 is a capsid-auxiliary protein that contains 8 CLRs centrally located in its sequence, and it is predicted to encode an α-helical C-terminus (Fig 7). The functional oligomeric state of gp12 is a trimer, assembled through the CLR region [69]. This was supported by circular dichroism analyses and confirmed by the cryo-electron microscopy reconstruction of the protein, which also indicated protein flexibility at one end [68,69]. Given the structural similarities with pUS10, it seems reasonable to assume that the functional form of pUS10 is also likely to be a trimer. Gp12 forms spikes that sit at the surface of the capsid, reversibly binding in a temperature-dependent manner, to the centre of the 60 capsid hexons. Binding of gp12 is thought to strengthen the adhesion properties of the virus [69]. Notably, in agreement with this, our clustering analysis, also predicts close functional association of pUS10 capsid proteins in HSV1.

**Fig 7.**
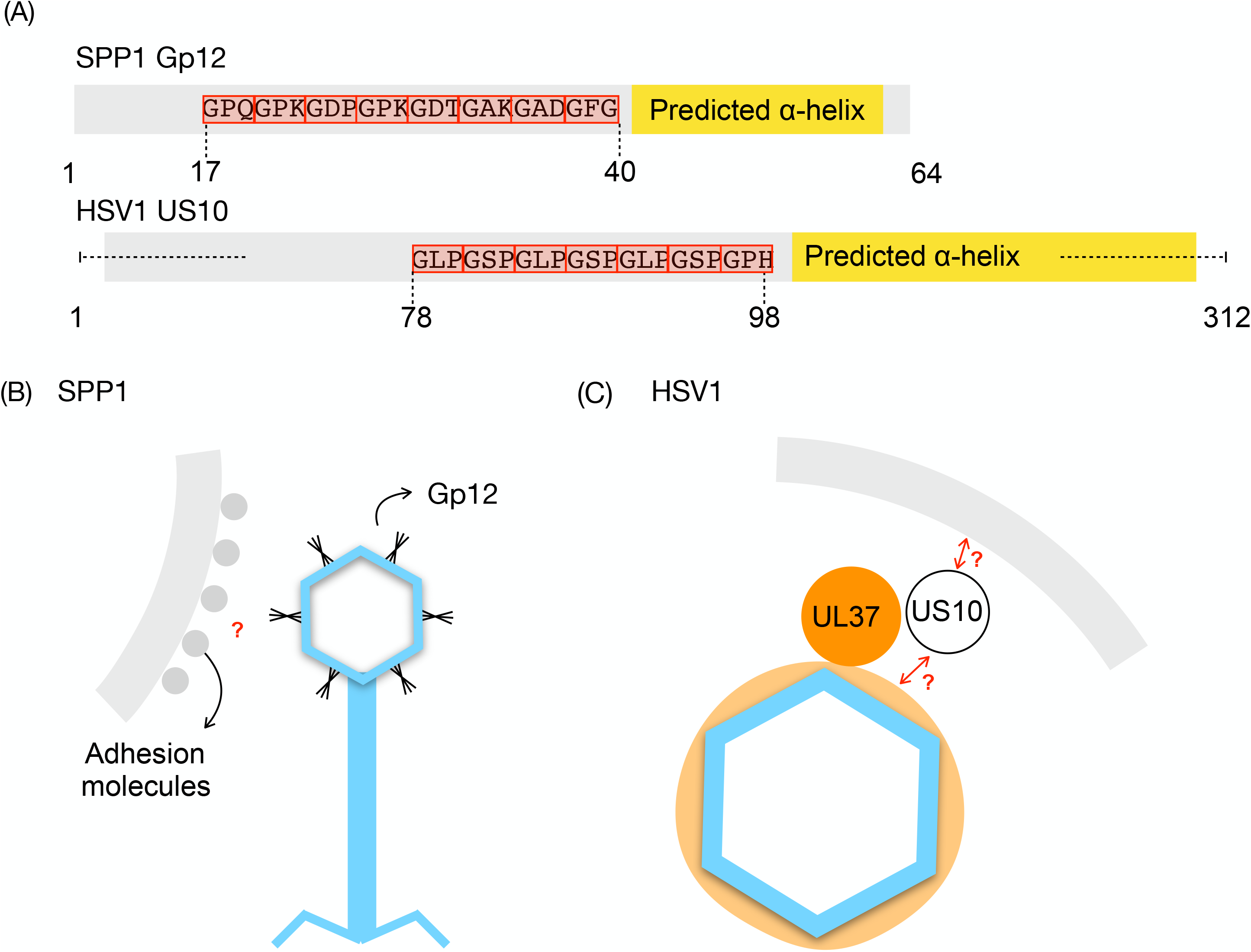
Structural and functional features of gp12 and pUS10. (A) Schematic comparison of the structural features of gp12 from bacteriophage SPP1 and pUS10 of HSV1. Protein sequences are shown in grey, CLRs as red boxes with the involved residues annotated, and predicted α-helical regions are highlighted as yellow boxes. (B, C) Comparison of the known biological function of gp12 (reversible binding to capsid hexons linked to phage surface adhesion, (B)) and that hypothesised for pUS10 (C). The interaction of pUS10 with HSV1 pUL37, predicted and experimentally supported in this study, would take place in sites of secondary envelopment, such as near Golgi- or trans-Golgi-network (TGN) derived membranes. Accordingly, pUS10 might facilitate the dynamic interactions with membranes in secondary envelopment of capsids through dynamic interactions with the latter, in a similar fashion to gp12’s interaction to adhesion surfaces, as well as potentially with membranes by means of its predicted transmembrane segment. Capsids (light blue) in (B) and (C) are drawn approximately to scale.

The identified evolutionary links between capsids of herpesviruses and tailed dsDNA bacteriophages have so far included the conserved HK97-core fold adopted by their major capsid proteins (MCPs), and their general assembly pathways (transitioning from an immature procapsid to a mature, stable, capsid) [68,71,72]. However, no evolutionary ancestry could be traced, based on either sequence or structural features, amongst capsid-associated or capsid auxiliary proteins of the two lineages. Here, we suggest that pUS10 could represent the first of such evolutionary vestiges. Clearly, SPP1 virions differ from those of herpesviruses in that they lack envelope, and, therefore, even in the presence of a common ancestor with gp12, one would expect pUS10 to have partially evolved new functionalities. In this context, the prediction of a transmembrane segment in the C-terminus of pUS10, which is not present in gp12, is interesting. Importantly, our network predicts an interaction between pUS10 and the inner tegument protein pUL37, which was further supported by our immunoaffinity experiments. The results of these experiments demonstrated a higher enrichment of pUS10 co-isolating with pUL37 later in infection (20 hpi). At these time points, pUL37 localises in secondary envelopment sites [37,38]. The functional role of an interaction between pUS10 and pUL37 could be that of facilitating the recruitment of the former to sites of secondary envelopment, to cooperatively function in subsequent capsid tegumentation events (Fig 7). An association of pUS10 to capsids via pUL37 would also explain why pUS10 has not yet been observed in the recent high-resolution structures of herpesviral capsids, [73–75]. On the basis of the structural similarities with gp12, it is reasonable to think that pUS10 could also exhibit reversible capsid/inner tegument-binding properties, similar to gp12, and establish dynamic interactions with capsid or capsid-associated proteins whilst, at the same time, interact with Golgi- or trans-Golgi-derived membranes, at the prospective capsids proximal pole of the forming virion, through its predicted transmembrane segment, and promote the final step in capsid trafficking during virion morphogenesis.

Together with pUS10, 9 out of the 14 proteins that our network data predict interact with pUL37, were co-purified with the latter in our immunoaffinity purification experiments (Fig 6D). Although these data cannot conclusively answer which binary interactions are direct ones among the co-isolated proteins, they add further support to the likely functional association of these proteins, and therefore add to the confidence of the PPI involved in the pUL37-centered subnetwork. Other interesting features of pUS10 will surely help defining its functionalities. For instance, the polyproline sequence located between two of the predicted C-terminus α-helices is also likely to be involved in PPI [76].

Although further investigations on the roles of pUS10 during viral infection are required, our analysis has revealed insightful new data on the structural features and potential evolutionary history of this intriguing and, so far understudied protein. Combined, our analyses have allowed to pose functional hypotheses that will further our understanding of the intricate tegumentation process of herpesvirus virions.

A second community sheds light to early tegumentation stages, specifically to the recruitment of a group of transcription and translation modulators, playing key roles at early stages of the lytic life cycle, into the virion particle. pUL48, also called VP16 or α-TIF, is a transcription activator that migrates with capsids to the host nucleus upon infection, and activates transcription of lytic genes [77]. The roles of pUL47 and pUL49 are more diverse, however both of them, together with pUL48 participate in regulating the activity of pUL41, also known as the virus host shut-off (VHS) protein, which inhibits protein synthesis [78]. Additionally, binary interactions among the four proteins seem to be required for their incorporation to the virion [39,40,79,80]. Yet, the stage at which this happens and the events that lead the process are unclear. Our results suggest that the recruitment of this module into the particle could be guided by the coordinated action of proteins pUL16, pUL11, pUL21, pUL14 and pUL56, potentially in peri- or juxtanuclear regions.

Two other communities are enriched in late stages of the tegumentation process and virion release. We observed a large number of proteins annotated with trafficking of TGN-derived vesicles and membrane-regulatory events. Particularly, our analysis suggests a strong clustering tendency between gK/UL20, pUL45, pUS2, and the poorly characterized α*-Herpesvirinae*-specific protein pUL55 [53]. Primary sequence analysis indicated pUL55 is a globular α/β protein with no predicted transmembrane regions. However, all four proteins gK, pUL20, pUL45, and pUS2, are membrane or membrane-associated proteins during intracellular stages of the virus life cycle, and gK, pUL20, and pUL45 are also envelope-associated proteins [54,81,82]. This finding supports the idea that pUL55 is a component of the outer most layer of the tegument, and its recruitment might be dependent on gK/UL20, pUL45, pUS2.

The outcomes of our protocol for predicting both binary and higher-order protein associations, bring novel insights and direction to future experimental testing. Further, we are currently working on the implementation of the introduced network assembly and analysis frameworks on other species of human herpesviruses. Widening this analysis to other members of this important group of human pathogens will also underscore both conserved and species-specific features of their interactome organization, and help to explain the observed phenotypes and evolution of their pathogenic strategies.

## Materials and Methods

### Data collection, curation, and integration

Binary PPI data from molecular interaction repositories were downloaded in the standardised MITAB 2.5 format [20] (Fig 1). Binary PPIs from PDB and EMDB entries were manually extracted. Only entries with assigned PubMed identifier were considered. In the case of EMDB entries, only those with resolutions ≤5Å at the time of this work were taken into account. When fitted atomic models were available, binary interactions between two proteins were extracted if the interface between them was larger than 500Å^2^ [83–85]. When fitted atomic models were not available, binary interactions were assigned as described in the primary citation. The resulting set of interactions were next manually annotated following the MITAB 2.5 format.

The data were collected for the species in S1 and S2 Tables. Only binary PPIs for which both interacting proteins were annotated with a taxonomic identifier corresponding to these species (all strains considered) were selected. Taxonomic identifiers were obtained from the National Center for Biotechnology Information (NCBI) [86] Taxonomy database (https://www.ncbi.nlm.nih.gov/taxonomy).

Protein identifiers were mapped, where possible, to UniProtKB [7] accession numbers. PPIs for which such mapping was not possible for one or both of the interacting proteins, were not further considered in our pipeline. The seven input PPI data sets were merged into a single non-redundant collection.

### Computational prediction of PPIs

PPI predictions were obtained using an interologues mapping approach [16]. This method (Fig 1D) predicts an interaction in species X if two proteins, known to interact in species Y, are conserved in both species X and Y. Here, protein conservation was assessed using sequence-based orthology predictions computed with the iterative Hidden Markov Model (HMM) profile comparison algorithm implemented by HHblits [17]. For each query sequence, the best match found in HSV1 satisfying all of the following conditions was considered a reliable putative homologue: ≥20% sequence identity, ≥30% sequence similarity, ≥50% query HMM profile coverage, and ≥95% probability of being a true positive. The resulting set of candidate homology relationships were used to infer interologues in each target interactomes.

### Integration of validated and predicted PPIs

Experimentally validated and computationally predicted interactions were merged into a single interactome data set. Strain redundancy was removed by mapping all protein sequences to reference strain accession numbers (HSV1 strain 17) using UniRef90 clusters.

### PPI scoring function

The scoring scheme integrated in our framework is inspired by the standardised MIscore function [18]. Under this scheme, PPIs that had experimental support in the target species (with or without additional support from computational predictions) were scored using the MIscore function. This was done through the MImerge service [18]. PPIs that did not have experimental support were scored with a new scoring function, defined as in Equation 1.

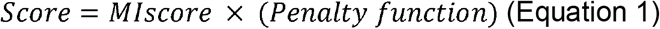

The new scoring function consists of first scoring an interaction using the MIscore [18], and next applying a penalty function to the returned score. This scaling factor (Equation 2) takes as reference the structure of the terms used in the MIscore function, but it redefines the meaning of their parameters to incorporate information on the number of species and prediction method used.

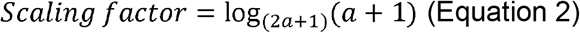

where

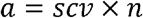

where *scv* is the score associated to the PPI prediction method (in this case interology mapping), and n is the number of different species from which the interaction was predicted.

The value of the penalty function increases asymptotically from ~0.5 to 1 with the number of species from which an interaction is predicted. Because we considered 10 orthologues to each target species, the values of the scaling factor in this study fall in the [0.5, 0.6] range. After applying the penalty function the values are normalised within [0, 1].

### Consensus clustering framework

Prior to clustering, the signal-to-noise ratio in the network data was increased as follows (Fig 3A). First, the subnetwork formed by proteins found in the mature virion particle of HSV1 (i.e. HSV1 virion subnetwork) was extracted from the full network. Next, self-interactions were removed as they do not add topological information about the network. Interactions between capsid and envelope proteins were also eliminated as they are precluded by the presence of the tegument in between the two former layers. Finally, the LCC was obtained and used as input to the consensus clustering framework.

A total of 14 different clustering algorithms, representing a broad spectrum of clustering techniques, were then applied to the LCC (Fig 3B), i.e. the K-means algorithm [87], agglomerative hierarchical clustering [88], K-means/hierarchical clustering [89], Fuzzy C-means [90], Model-based clustering [91,92], Markov Cluster algorithm (MCL) algorithm [93], Density-based clustering [94], Edge betweenness [95,96], Louvain method [97], and Leading eigenvector [98], Fast greedy [28], Walktrap [99], and InfoMap [100] algorithms. Where needed, partition selection and parameter optimisation were conducted based on modularity [95] maximisation. The clustering results of those partitions yielding positive modularity were integrated into a consensus agreement matrix (CAM) (Fig 3C and S4 Table). The condition of positive modularity discarded partitions were each and every node was singled out as a cluster by itself, the entire network was treated as a super-cluster, or the graph under such partition had less local structure than random models. The columns and rows of the consensus agreement matrix represented the network proteins; the matrix values indicated the fraction of accepted base partitions classifying two proteins in the same cluster.

A final consensus partition was derived from the CAM by applying a filtering threshold (Fig 3D and S5 Table). For thresholds between 0 and 1, by increases of 0.5, we iteratively calculated a consensus partition in the following way. CAM cells with values equal to or below the threshold were ignored; the remaining cells defined a partition for which the modularity was calculated. At the end of this iterative process, the partition associated to the highest modularity was chosen as consensus.

### Sequence analysis

The canonical sequence of pUS10 and pUL55 in the HSV1 reference proteome were obtained from UniProtKB [7]. For each sequence, the following analysis was conducted. ScanProsite [101] was used to scan the sequences for sequence motifs. Potential sequence homologues within the entire UniProtKB database [7], were searched for using HHblits [17]. Next, a consensus prediction of secondary structure elements was inferred from the results of four different secondary structure prediction tools, i.e. SPIDER^2^ [102], PSIPRED [103], JPred4 [104], and PSSpred [105]. Probabilities associated to the returned predictions were not integrated in the consensus analysis. Similarly, consensus predictions for transmembrane segments were derived from five different algorithms, i.e. Dense Alignment Surface (DAS) [106], Phobius [107], PHDhtml [108], TMpred [109], and MEMSAT-SVM [110].

From TMpred predictions only significant regions (defined as regions with score above 500) and core residues were taken into consideration. From Phobius, only residues with probability of belonging to a transmembrane region above 0.1 were considered. Finally, a consensus prediction for disordered regions was built from the results of two algorithms, i.e. DISOPRED [111] and MetaDisorder [112].

### Infection of human fibroblasts with HSV1 and live cell imaging

We used the HSV1(17^+^)Lox-UL37GFP strain, a generous gift from B. Sodeik and previously characterized in [37], here denoted HSV1-UL37EGFP, and as a control, the HSV1(17^+^)Lox-P_MCMV_GFP strain, denoted HSV1-EGFP, which expresses EGFP alone inserted between the pUL55 and pUL56 ORFs, under the control of the murine cytomegalovirus promoter [113]. Viruses were propagated, isolated, and titered in Vero cells (ATCC CCL81) grown in DMEM containing 10% FBS and 1% penicillin/streptomycin (P/S), as previously described [8]. Primary human foreskin fibroblast cells were infected with HSV-1 strains at 10 plaque forming units/cell using a cold-synchronized protocol [114]. The progression of infection was visualized by live-cell imaging on a Nikon Ti-Eclipse epifluorescence inverted microscope from 2 hpi to 24 hpi. Images were viewed and analysed by ImageJ [115].

### Immunoaffinity Purification Quantitative Mass Spectrometry

For IP-MS experiments, cells were infected as above with HSV1-UL37GFP or control HSV1-GFP, in duplicate. Infected cells were collected at 8 and 20 hpi (HSV1-UL37GFP) or 20 hpi (HSV1-GFP) in ice-cold PBS and pelleted by centrifugation (~1 x 10^7^ per time point per replicate). Cell pellets were washed in ice-cold PBS and lysed hypotonically. Cytosolic lysates were adjusted to 20 mM HEPES-KOH, pH 7.4, containing 0.11 M potassium acetate, 2 mM MgCl_2_, 0.1% Tween 20, 1 μM ZnCl_2_, 1 μM CaCl_2_, 250 mM NaCl, and 0.5% NP-40, mixed by Polytron homogenization, and centrifuged at 8,000 x g for 10 min at 4°C. The supernatant was recovered and subjected to immunoaffinity purification using magnetic beads conjugated with in-house generated rabbit anti-GFP antibodies, as previously described [114,116].

Immunoisolated proteins were processed by a Filter-Aided Sample Preparation method using Amicon ultrafiltration devices (Millipore, 30 kDa MWCO) as described [117], except 0.1 M Tris-HCl, pH 7.9 was replaced with 0.1 M triethylammonium bicarbonate (TEAB). Following overnight trypsin digestion and clean-up, peptides (4 μl) were analyzed by nanoliquid chromatography-tandem mass spectrometry on a Dionex Ultimate 3000 RSLC coupled directly to a LTQ Orbitrap Velos ETD configured with a Nanospray ion source (Thermofisher Scientific).

The Proteome Discoverer software (ver. 2.2) was used for post-acquisition mass recalibration of precursor and fragment ions masses, MS/MS spectrum extraction, peptide spectrum matching and validation, calculation of TMT reporter ion intensities, and assembly of quantified into protein groups. Protein groups and TMT protein abundances for herpesvirus proteins with a minimum of 2 unique quantified peptides were exported to Excel. IP protein enrichment ratios for each time point and replicate were calculated as the TMT abundance ratio of pUL37GFP / GFP. Proteins with IP enrichment ratios of ≥ 2-fold in at least one time point in both replicates were considered specific associations. The TMT abundance ratio for proteins in the 20 vs 8 hpi pUL37GFP IPs were calculated after normalization by the pUL37 TMT abundance. Further details on data collection and analysis can be found in S1 Text.

## Acknowledgments

We are grateful for funding from Human Frontiers Science Program (RGY0079/2009-C) to K.G., M.T., and I.M.C., from the NIH (GM114141) and a Mallinckrodt Scholar Award to I.M.C., from the Wellcome Trust (209250/Z/17/Z) to M.T. and K.G., and (107806/Z/15/Z) and a WT Core Award (203141/Z/16/Z) to K.G., and from the MRC (MR/M019292/1) to M.T. and K.G.

## Supporting Information

**S1 Text. Details on Immunoaffinity Purification Quantitative Mass Spectrometry experiments.**

**S1 Fig.**
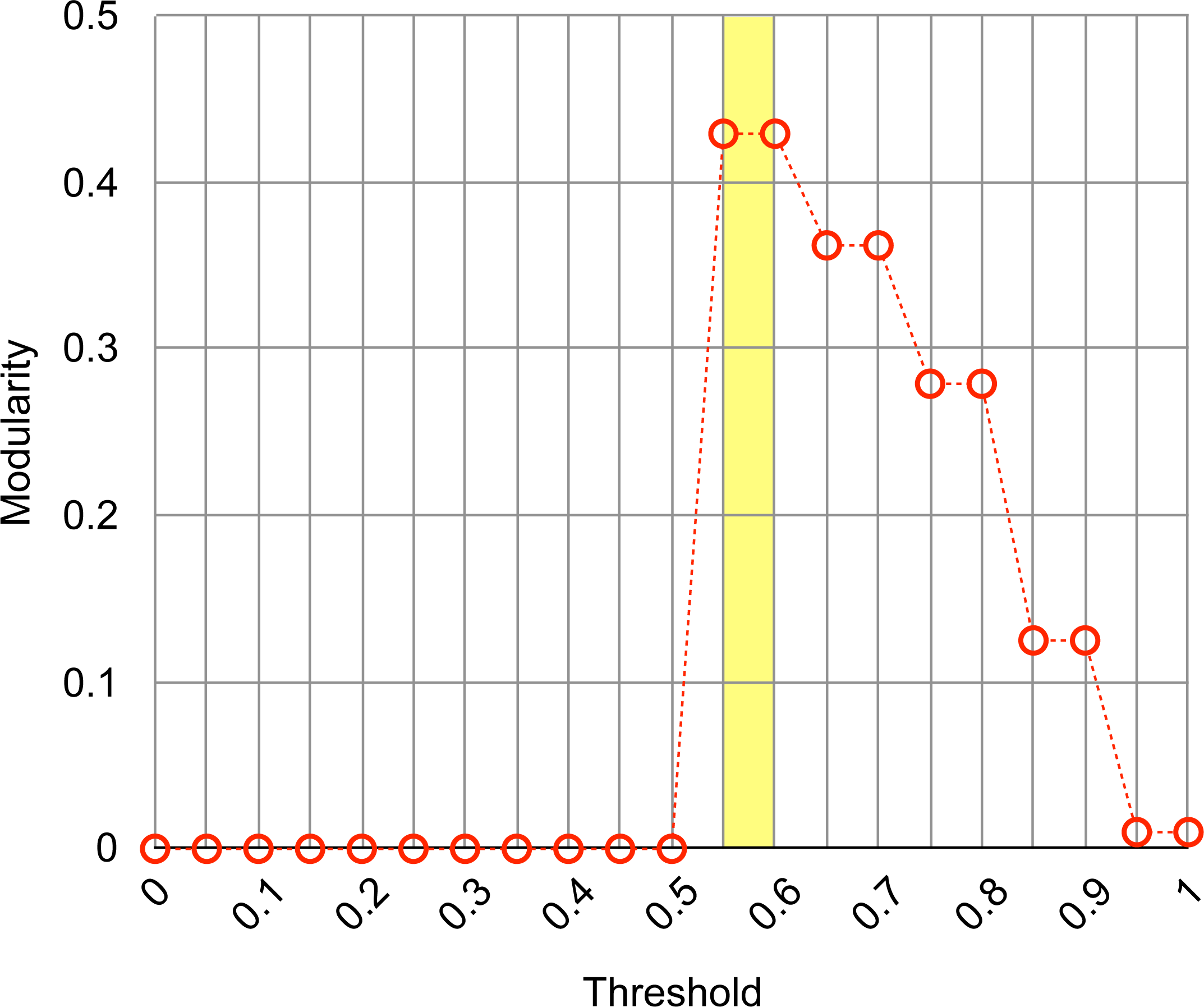
Modularity values of the consensus partition as a function of the filtering threshold. Modularity values achieved by each of the partitions derived from the consensus agreement matrix, by filtering values below the specified thresholds (ranging from 0 to 1 in increments of 0.5). The dashed line is only shown to emphasise the trend of the modularity values. The yellow-shaded area of the plot highlights the threshold range for which the highest modularity was achieved.

**S2 Fig.**
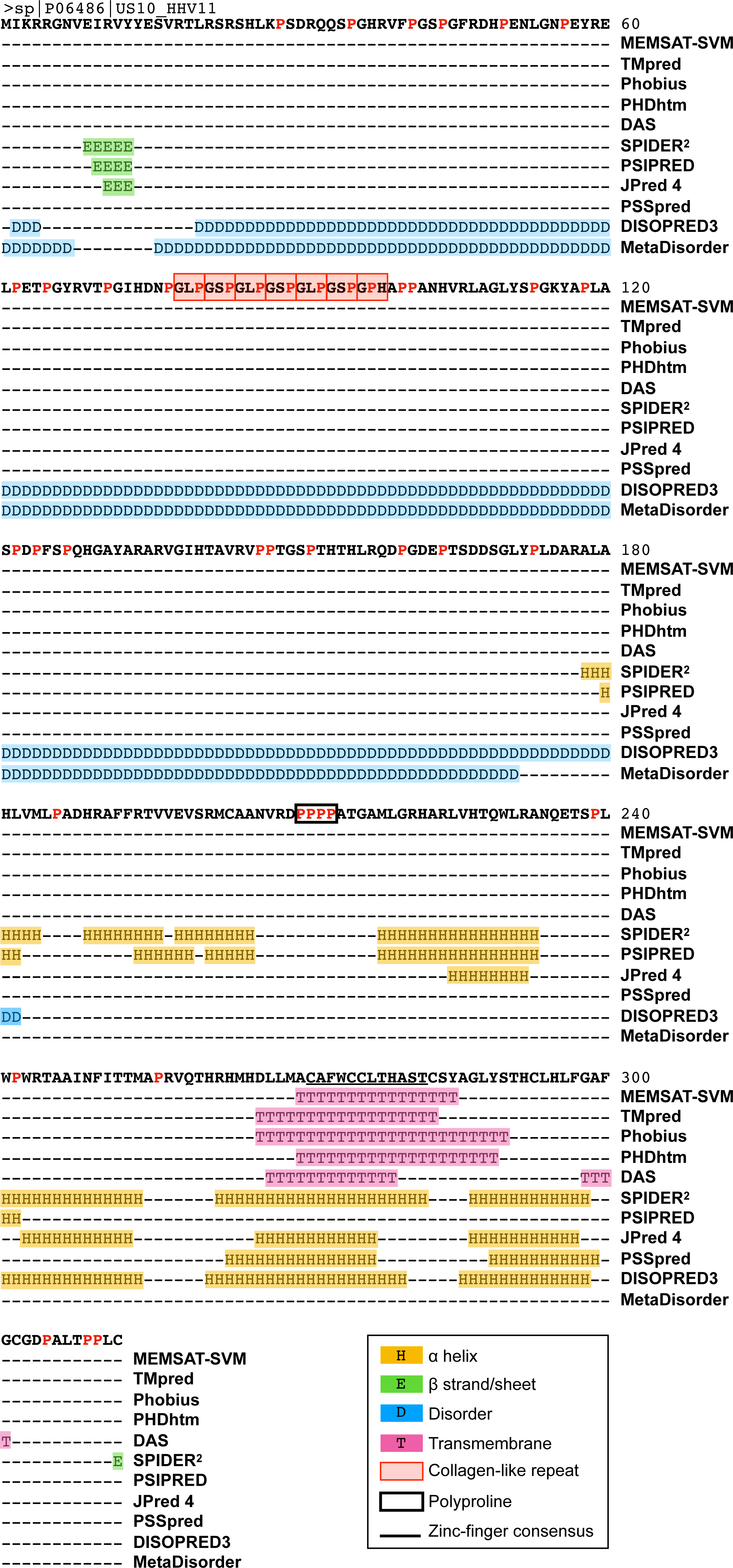
Primary sequence analysis of US10. Previously reported and newly identified features are indicated. Predictions from each software tool are shown. Predicted disordered regions, α-helices and transmembrane helices are indicated in blue, yellow and pink, respectively. The identified CLRs are shown in red boxes. Individual prolines are highlighted in red. The four-residues polyproline sequence is indicated with a black box. The previously identified consensus zinc finger sequence [33] is underscored. The final assignment of the secondary structure elements was based on the consensus of individual methods (prediction confidence scores were not taken into account).

**S3 Fig.**
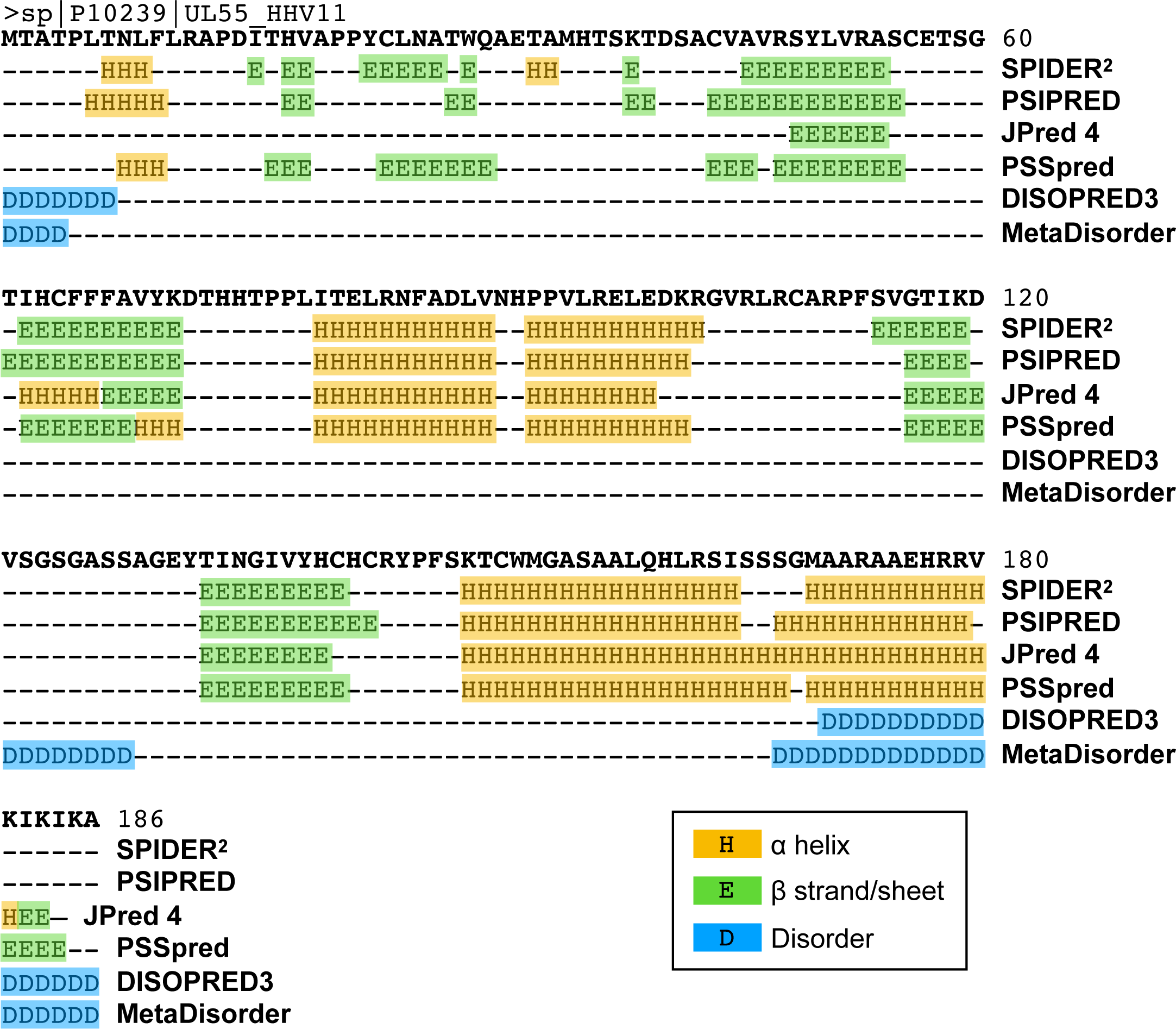
Primary sequence analysis of UL55. Predictions from each software tool are shown. Predicted disordered regions, α-helices and β-strands are indicated in blue, yellow and green, respectively. The final assignment of the secondary structure elements was based on the consensus of individual methods (prediction confidence scores were not taken into account).

**S1 Table. Herpesvirus species for which PPI data were collected as input for the PPI network assembly framework.**

**S2 Table. Taxonomic identifiers associated to species in S1 Table and used to extract PPIs from input resources.**

**S3 Table. Protein-protein interaction network reconstructed for HSV1.**

For each interaction, the interacting proteins, detection methods, associated PubMed IDs, types of interaction, confidence score, and whether the interaction was computationally predicted and/or experimentally supported, is indicated.

**S4 Table. S4 Table. Parameters associated to the partitions returned by the 14 clustering algorithms.**

**S5 Table. Modularity values associated to the partitions derived from the consensus agreement matrix, at different thresholds.**

**S6 Table. Proteins co-purifying with pUL37 by immunoaffinity purification mass-spectrometry.**

## Abbreviations

CAM: consensus agreement matrix
CLR: collagen-like repeat
DIP: Database of Interacting Proteins
dsDNA: double-stranded DNA
EBV: Epstein-Barr virus
EGFP: enhanced green fluorescent protein
EMDB: Electron Microscopy Data Bank
ePPI: experimentally detected PPI
GO: gene ontology
HCMV: human cytomegalovirus
HMM: Hidden Markov Model
HPI: hours post infection
HSV1: herpes simplex virus type 1
IP: immunoaffinity purification
LCC: largest connected component
MCP: major capsid protein
MS: mass spectrometry
PDB: Protein Data Bank
PPI: protein-protein interaction
oPPI: PPI in orthologous species
pPPI: computationally predicted PPI
TGN: trans-Golgi network
TMT: tandem mass tags
VZV: varicella-zoster virus

